# Universal super-resolution for subcellular fluorescence imaging

**DOI:** 10.1101/2025.10.26.683961

**Authors:** Xiangcong Xu, Renlong Zhang, Qinglin Chen, Xinwei Gao, Zhao Xie, Danying Lin, Wei Yan, Xuehua Wang, Leiting Pan, Liwei Liu, Jia Li, Junle Qu

## Abstract

Super-resolution fluorescence microscopy has emerged as an essential instrument for investigating subcellular structures and monitoring intracellular physiology. Despite rapid advances driven by deep learning, most existing approaches are modality specific and exhibit limited generalizability, restricting their practical applicability. Here we introduce universal super-resolution (UniSR), a framework that operates across microscopes and subcellular structures without optical modifications or prior sample knowledge. UniSR is initially pre-trained on simulated image pairs to establish mappings from low-to high-resolution features and subsequently fine-tuned with only one experimental image pair, reducing data demand and acquisition time. Across wide-field and point-scanning modalities, UniSR enhances resolution to match structured illumination, single-molecule localization, or stimulated emission depletion microscopy. Applied to diverse biological samples, it reveals nanoscale structures, dynamic processes, and organelle interactions with high spatiotemporal fidelity. UniSR provides a practical and broadly generalizable tool for super-resolution imaging in live-cell research.

## Introduction

Super-resolution (SR) fluorescence microscopy provides unparalleled access to the nanoscale spatial details of cellular structures and dynamic biological processes, thus facilitating the exploration and in-depth investigation of fundamental life activities^1–3^. Over the past two decades, major advancements have been made in the development and optimization of SR fluorescence microscopy techniques. These encompass structured illumination microscopy (SIM)^4, 5^, single-molecule localization microscopy (SMLM)^6, 7^, and stimulated emission depletion (STED) microscopy^8, 9^. Each of these techniques utilizes distinct physical principles to surpass the diffraction limit.

However, the improvement in spatial resolution is inherently accompanied by trade-offs in other imaging conditions^10^, including exposure time, temporal resolution, and illumination intensity, which are also crucial for accurately resolving subcellular structures and capturing dynamic biological processes^11^. This has consequently sparked considerable interest in computational SR methods aimed at enhancing the resolution of diffraction-limited images without significantly compromising other important metrics. In particular, recent breakthroughs in deep learning have exhibited strong ability to directly transform low-resolution (LR) images into SR counterparts without any alterations to optical systems^12–18^.

Despite their achievements, contemporary deep-learning-based SR methods encounter two primary limitations that impede wider applications. Firstly, the predominant models are modality-specific and exhibit poor generalization. When confronted with substantial domain disparities between datasets, the performance of these models deteriorates significantly. Owing to the extreme diversity of subcellular biological structures and the limited representational capacity of deep neural networks, existing methods frequently require the training of a dedicated model for each specific specimen to attain reliable results^19^. Secondly, given the data-driven nature of deep learning, the performance of these models is highly contingent upon the quality and quantity of the training data. A large number of high-quality SR images are required to serve as training targets. This elevates the implementation complexity of deep-learning-based SR methods^20^, as the experimental acquisition of matched LR-SR training image pairs is experimentally demanding, especially in cases of rapid dynamics or severe photobleaching in biological samples.

To tackle these challenges, we propose a universal super-resolution (UniSR) model, a transfer-learning-based^21–23^ universal SR approach designed to be broadly applicable, user-friendly, and data-efficient. This model is pre-trained with simulated data and subsequently transferred to diverse tasks. Through fine-tuning with only one image pair instead of training from scratch on specific datasets, it achieves substantial performance improvements. It enhances the resolution of raw images captured by both wide-field (WF) fluorescence and point-scanning microscopes, enabling SR imaging of 2D/3D organelle interactions, cytoskeletal and organellar dynamics, and diverse subcellular structures across different cell types, such as COS-7, Beas2B, HeLa, U2OS, Vero, and mouse embryonic fibroblasts. By decoupling the need for extensive experimental training data, UniSR provides a powerful and practical computational tool that makes SR imaging accessible to users in a variety of bioimaging applications, paving the way for further discoveries in live-cell research.

## Results

### Development and characterization of UniSR

To attain resolution enhancement across both modalities (WF and point scanning microscopy) and cellular structure (diverse organelles), this paper proposes a transfer-learning-based method for super-resolving LR images. Figure 1 illustrates the workflow of our approach, UniSR, and its training can be partitioned into two stages (Figure 1B). In the first stage, the model is pre-trained on an adequate amount of simulated data of linear structures (Methods), learning fundamental visual features such as edges and the basic mapping between LR and SR images. Subsequently, in the second stage, the pre-trained model is fine-tuned using only one image pair, enabling rapid adaptation to cellular structures of different morphologies and alternative imaging modalities. Our model is tested on a wide range of subcellular structures (Figure 1A), including microtubules (MTs), lysosomes, and outer mitochondrial membranes (OMM), which are three of the most prevalent biological structures in live-cell experiments; F-actin filaments and inner mitochondrial membranes (IMM), which exhibit the highest structural complexity, as well as clathrin-coated pits (CCPs), endoplasmic reticulum (ER), nuclear pore complexes (NPCs) and Golgi apparatus.

**Figure 1.**
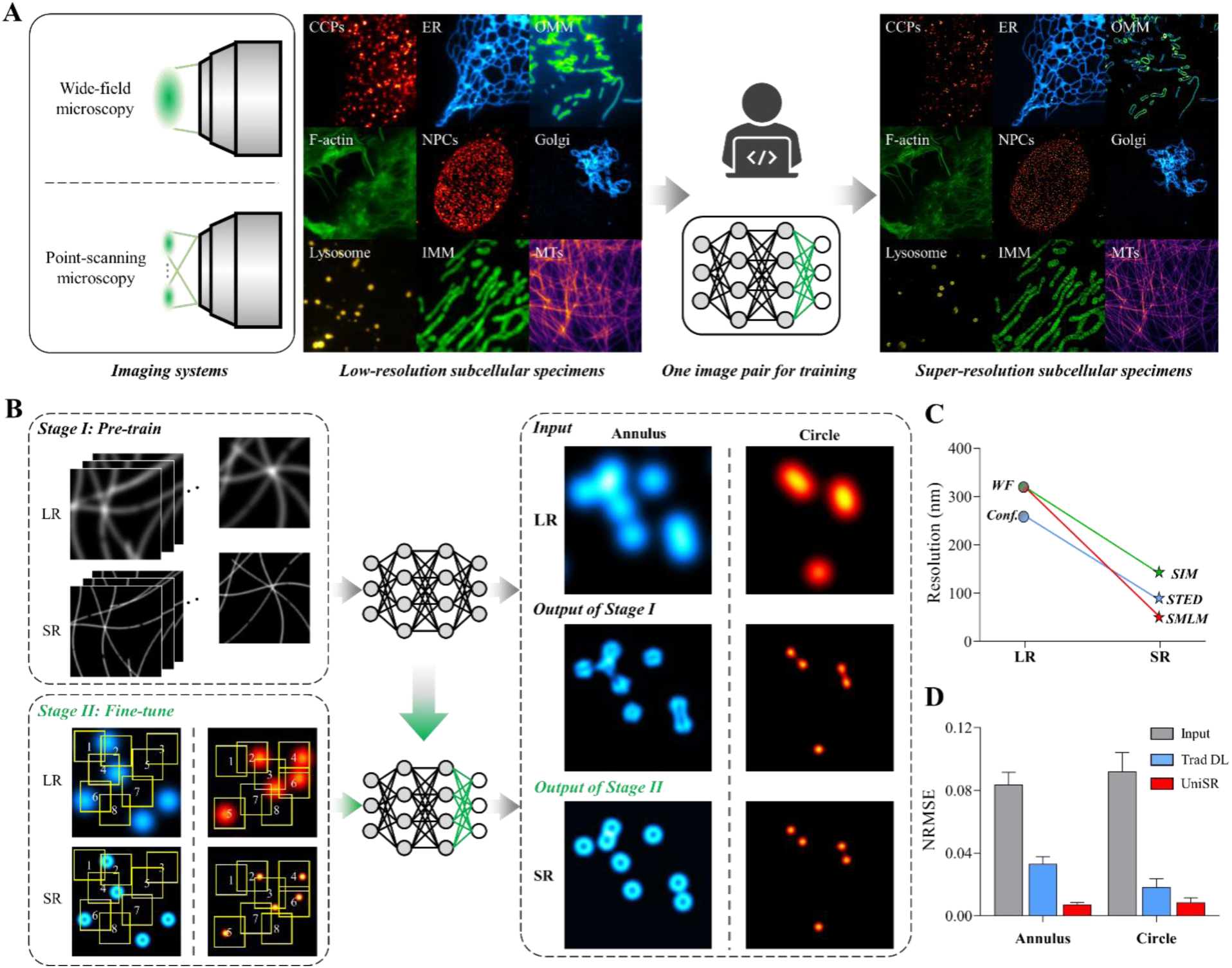
Principle of UniSR. **A** Overview of the UniSR workflow. **B** Schematic diagram of the training mechanism of UniSR. **C,D** Quantitative metrics across WF-to-SIM, WF-to-SMLM, and confocal-to-STED transformations.

During the fine-tuning phase, we freeze the majority of convolutional layers to retain their pre-learned weights, and only update the weights of the final convolutional layer. This strategy capitalizes on the generalizability of early-layer features for visual processing, while enabling task-specific adaptation in the later layers that integrate these foundational features. This is especially critical for our domain-specific data^24^. A relatively low learning rate is utilized during fine-tuning to preserve the learned useful representations and guarantee training stability. Considering the utilization of only one image pair, this approach optimizes the extraction of feature information from limited data.

We propose a novel multi-window synchronized sampling strategy (Methods). Eight sampling windows are randomly distributed across the image to capture regions of interest (ROIs). Each window undergoes independent data augmentation (rotation, flipping, or scaling) prior to being stacked for model training. These randomly positioned windows cover diverse regions of the image, capturing multi-perspective feature representations. This strategy enhances the model’s comprehension of structural variations, improving SR performance and enhancing adaptability to different imaging modalities and cellular structures.

To verify the necessity and efficacy of both pre-training and fine-tuning, the proposed UniSR is applied to simulated LR-SR data, and the results are presented in Figure S1. The first column showcases the LR input images along with their corresponding ground truth (GT). It can be noted that the SR images predicted by the models without fine-tuning (the third and fifth columns) demonstrate significantly higher normalized root mean square error (NRMSE) compared to those generated by fine-tuned models (the second and fourth columns). Similarly, the SR images predicted by the model without pre-training (the sixth column) also display significantly higher NRMSE values, even though it is fine-tuned with data that is structurally identical to the test data. The model without pre-training (the sixth column) is only capable of reducing the size of the circles but fails to resolve two adjacent circles.

As depicted in Figure S1B, models incorporating both stages consistently exhibit lower NRMSE, higher peak signal-to-noise ratio (PSNR)^25^, and superior structural similarity index (SSIM)^26^ values, which indicate that the outputs are closer to the GT images. These image fidelity metrics for the inferred SR images confirm that both pre-training and fine-tuning are crucial for enhancing image resolution. This conclusion is further corroborated by t-distributed stochastic neighbour embedding (t-SNE) analysis^27^ in Figure S1C. The feature distributions from models pre-trained on identical structural data types cluster closely. In contrast, models without pre-training exhibit dispersed and chaotic feature distributions. The validation is repeated using 1000 simulated image pairs of annular, circular and linear structures. It is consistently observed that models lacking either pre-training or fine-tuning perform inferiorly compared to transferred models.

Furthermore, it is discovered that structural complexity^15^ exerts an influence on model performance. As depicted in Figure S2, models pre-trained on linear structures exhibited higher NRMSE values and lower SSIM and PSNR values. This indicates that linear structures are more complex, thus posing the greatest challenge for SR reconstruction tasks. Therefore, the model pre-trained on large-scale linear datasets is selected as the optimal configuration.

### UniSR-enabled super-resolution imaging: resolution enhancement from wide-field to SIM-level images

We commence by presenting the resolution enhancement capacity of the proposed UniSR for WF images. Although WF microscopy is proficient in capturing dynamic biological processes owing to its high imaging speed, its spatial resolution is still restricted. To validate its performance across a wide range of subcellular structures, we utilize established public datasets focusing on CCPs, ER, MTs, F-actin filaments, lysosomes, and OMM^14, 15^.

Figure 2A illustrates that UniSR expands the limited spatial frequency bandwidth of WF microscopy to be consistent with that of SIM, without the necessity of multi-frame patterned illumination. In Figures. 2B and 2C, UniSR takes WF images (the first column) as input and generates SR images (the second column), while the corresponding SIM image (the fourth column) serves as the GT for comparison. When compared to the input WF images, the network outputs reveal structures finer than the diffraction-limited scale. For instance, the annular structure of CCPs and the densely interwoven network of MTs or F-actin filaments can be observed. Some structures are too closely spaced to be differentiated in the original WF image, yet our method successfully resolves them and exhibits a good correspondence with the SIM images of the same sample regions, as evidenced by the intensity profiles along the white dashed lines.

**Figure 2.**
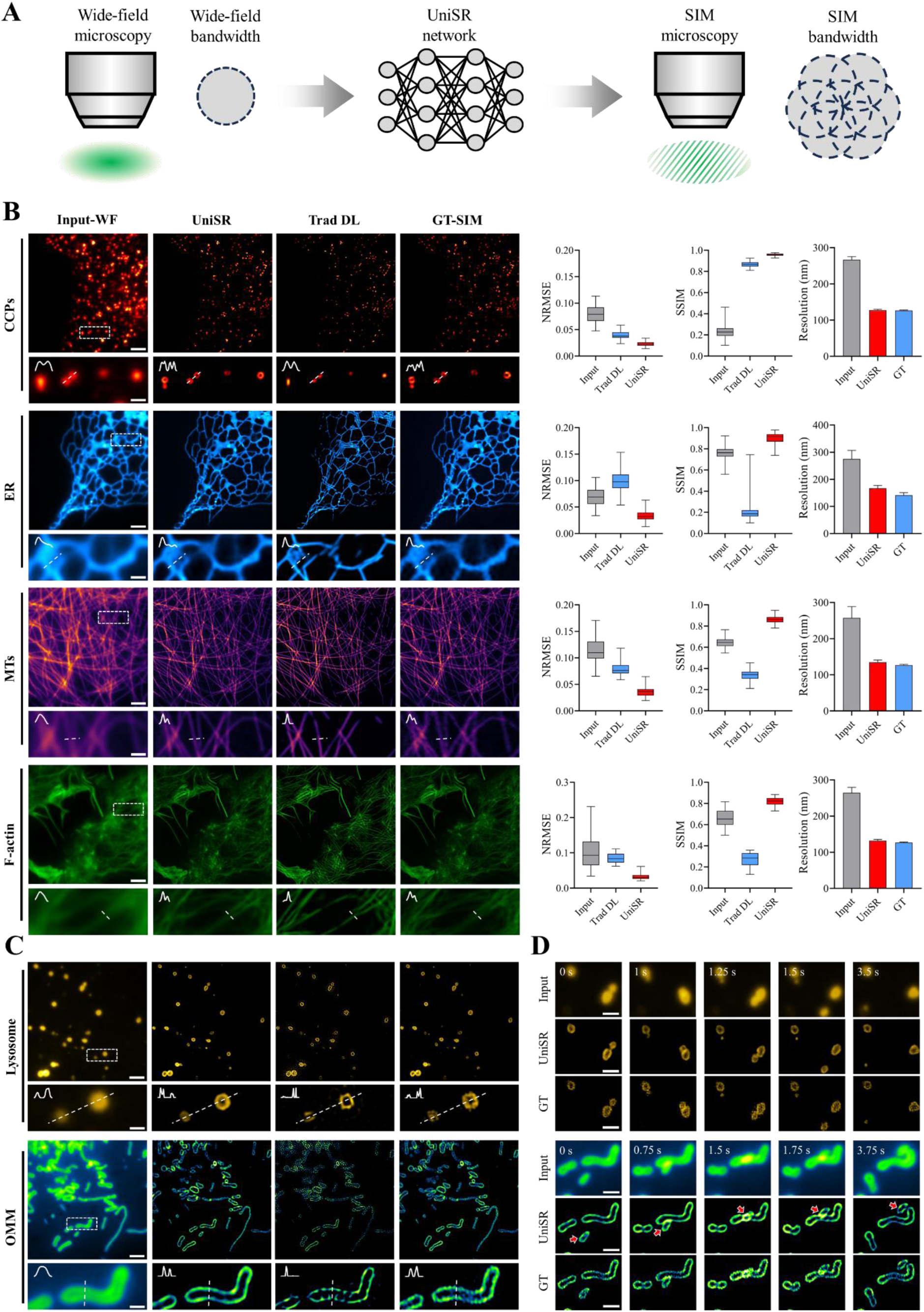
UniSR achieves SIM-level resolution from WF images. **A** Schematic illustration of resolution enhancement from WF to SIM via UniSR. **B, C** In the first column, diffraction-limited WF images are presented as the input to the network. The second and third columns display the SR images inferred by UniSR and Trad DL, respectively. The fourth column shows the corresponding SIM images of the same ROI. Magnified views of white boxed regions are shown below. Quantitative comparison of NRMSE, SSIM and resolution for input WF images, inferred images generated by UniSR and Trad DL, and the GTs. **D** Magnified time-lapse images captured at different time points. Scale bars: 5 μm (**B**-**D**), 1 μm (zoom-in regions in **B**-**D**).

We also contrast the outcomes of UniSR with those of a traditional deep learning (Trad DL, the third column) model. The Trad DL model shares the same network architecture and is trained from scratch without fine-tuning. Although the output of Trad DL yields marginally sharper features, this is achieved at the expense of losing structural information. In contrast, our method achieves superior resolution while preserving high-fidelity, demonstrating excellent consistency with the GT images.

It is worth emphasizing that for each biological structure, regardless of its intricacy, we fine-tune the pre-trained model using only one pair of WF-SIM data. To quantify the improvement in image quality by UniSR, we evaluate the SR outputs using three quantitative metrics: NRMSE, SSIM, and resolution^28^. These measurements (Figure 2B) confirm that the proposed method consistently enhances the resolution across various subcellular structures, highlighting its broad applicability to WF microscopy.

We observe that UniSR elevates the resolution of WF images to a level comparable to that of SIM, as the upper resolution limit of the network is determined by the SIM data employed for fine-tuning. To examine the applicability of UniSR to live-cell imaging, we investigate the dynamics of lysosomes and OMM through time-lapse imaging. The enhancement in spatial resolution facilitates the visualization of mitochondria in live cells (Figure 2D). As indicated by the red arrows, two mitochondria undergo physical contact and subsequent separation without fusion, which are not clearly discernible in the LR time-lapse images.

A significant advantage of our method lies in its robustness against noise. To systematically assess the robustness of UniSR to different degrees of experimental noise, we employ the dataset under ultra-high (U), high (H), medium (M) and low (L) SNR levels. In comparison with GT-SIM images, the proposed method effectively discriminates biological features from artifacts, while maintaining stable NRMSE values across various noise conditions and subcellular structures (Figure S3). This method achieves an optimal balance between feature preservation and noise suppression. UniSR outputs more refined details even in regions characterized by severe noise and background fluorescence, demonstrating its efficacy in generating high-fidelity SR images from low-SNR WF inputs.

Another substantial advantage of UniSR is its consistent resolution enhancement across a diverse range of FOV scales (from small to large), without any significant deterioration in structural accuracy. This stability is presented in Figures. S4-S6, where images with different scale bars represent distinct FOVs. A prepared four-color slide (CSR Biotech, Guangzhou, China) is imaged using a custom-built WF microscope based on an inverted fluorescence microscope (Ti2-E, Nikon, Japan). The system is equipped with an sCMOS camera (SD1000C/M, Image Technology, Suzhou, China) and three objectives: a 20× air objective (Nikon ELWD S Plan Fluor, NA 0.45); a 40× air objective (Nikon ELWD S Plan Fluor, NA 0.6); and a 60× oil-immersion objective (Nikon APO, NA 1.4). To enable the direct processing of large-scale images without computationally intensive sliding window and stitching approaches, we simplify the UniSR architecture to a more efficient, conventional U-Net. UniSR enhances the resolution of multi-scale WF images acquired with 20×/0.4 NA, 40×/0.6 NA, and 60×/1.4 NA objectives, substantially improves the visualization of finer subcellular structures such as MTs, mitochondria, and F-actin filaments, while maintaining spatial integrity across large imaging areas. A large FOV captures the whole-cell morphology and spatial distributions, whereas a small FOV resolves cellular details. Our method maintains consistent performance when processing these cross-scale datasets, effectively enhancing the image resolution and quality.

All these results illustrate that UniSR represents a robust methodology capable of elevating the resolution of WF images to SIM levels, presenting considerable potential for the rapid and high-fidelity imaging of subcellular organelles.

Moreover, we investigated two factors influencing network prediction performance: the type of pre-training data and the sizes of the fine-tuning dataset. In Figure S7, a comparison is made between the performance of UniSR pre-trained on simulated data (the second column) and that pre-trained on experimentally acquired microtubule data (the third column). For three representative cellular structures, namely CCPs, ER, and F-actin, the former achieves performance comparable to the latter. Error maps (the fourth column), which depict pixel-wise intensity differences between the outputs of these two pre-trained models, disclose nearly identical overall morphologies. This approach significantly curtails data acquisition costs while upholding the quality of SR images. It is especially advantageous for imaging modalities in which the acquisition of a substantial amount of high-quality experimental data remains a challenge. More crucially, this advancement substantially reduces the dependence of deep-learning-based microscopy on experimentally acquired training data.

In Figure S8, when the quantity of fine-tuning data is increased from 1 to 30 frames, only a marginal reduction in NRMSE is observed. In essence, the expansion of the fine-tuning data pool exerts minimal influence on the network performance. This aligns with the expectations of efficient learning during fine-tuning, as UniSR extracts discriminative features from only one image pair, effectively exploiting the maximum information from available data. This indicates its applicability in data-scarce scenarios where the acquisition of sufficient training datasets is technically arduous due to inherent biological rarity (e.g., rare cellular subtypes) or imaging limitations (e.g., low-abundance subcellular structures).

### UniSR-enabled super-resolution imaging: resolution enhancement from wide-field to SMLM-level images

Subsequently, we conduct fine-tuning of UniSR with one pair of WF-SMLM images to further elevate the resolution of WF images to match that of SMLM. Figure 3A demonstrates that UniSR directly generates an SR image from a single WF frame, obviating the need for sequential localization of individual fluorescent molecules and reconstruction from thousands of frames required for SMLM. This method is tested on the WF microtubule data (Methods)^29, 30^ and a public available experimental dataset ^16^. As depicted in Figure 3B, UniSR effectively resolves dense and intricate microtubules, producing sharper structures that are comparable to the reconstructions obtained through conventional SMLM from thousands of raw images. In contrast, Trad DL approaches generate a multitude of artifacts and fail to yield valid reconstruction results. It is noteworthy that the achieved resolution reaches the SMLM level (50 nm), as the maximum performance of the network is determined by the resolution of the fine-tuning SMLM data. Our proposed method also exhibits robustness against variations in imaging conditions, such as different spectra (Figure 3C), by successfully converting the WF images of microtubules immunostained with Alexa Fluor 488, 568, and 647 into their SR counterparts^16^.

**Figure 3.**
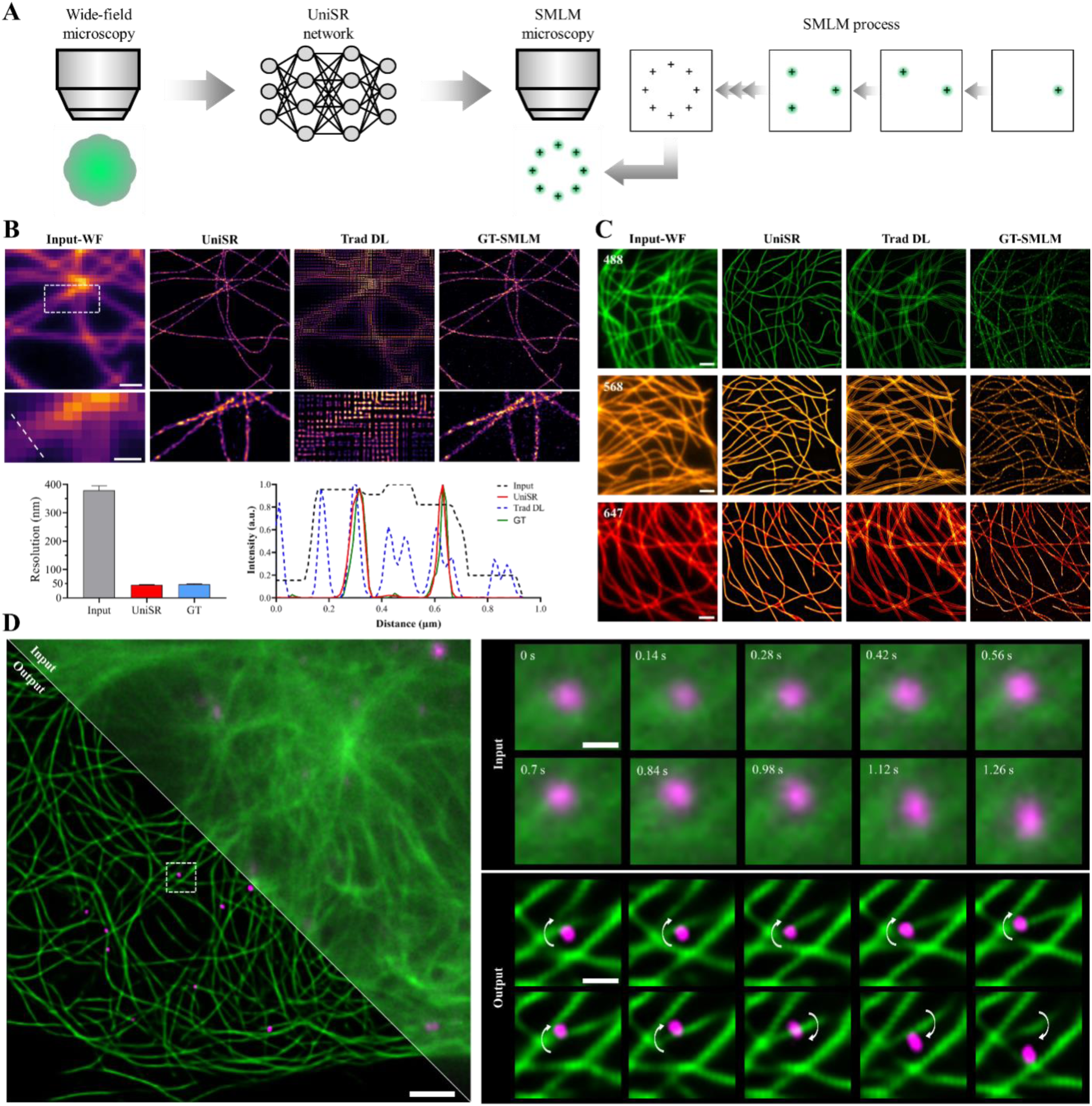
UniSR achieves SMLM-level resolution from WF images. **A** Schematic representation of resolution enhancement from WF to SMLM through UniSR. **B** The first column consists of diffraction-limited WF images utilized as input for the network. The second and third columns present the inferred SR images generated using UniSR and Trad DL, respectively. The fourth column displays the corresponding SMLM images of the same ROI. Magnified views of the regions enclosed by the white dashed boxes are presented below. Comparison of the resolution among the input WF images, UniSR outputs, and the GTs. Comparison of the intensity profiles along the white dashed lines. **C** The first column comprises input WF images obtained from microtubules immunostained with Alexa Fluor 488, 568, and 647 separately. The second and third columns showcase the inferred SR images produced using UniSR and Trad DL, respectively. The fourth column provides the corresponding SMLM images of the same ROI. **D** The first column contains the input WF image (top right) and the corresponding UniSR output (bottom left). The second column shows magnified views of the regions enclosed by the white dashed boxes at different time points. Scale bars: 1 μm (**B**-**D**), 0.5 μm (zoom-in regions in **B**), 0.2 μm (zoom-in regions in **D**).

The dynamic interplay between MTs and vesicles underpins intracellular transport and signaling, both of which are vital for maintaining cellular homeostasis. This rapid interaction requires imaging with high spatiotemporal resolution^32^. Leveraging the capabilities of UniSR, we concurrently monitor the dynamics of vesicles and microtubules. Over time, the vesicle moves along a quasi-circular trajectory around the microtubule, as indicated by the white arrows in Figure 3D. This finding is consistent with the results of a previous study^16^. These subtle interaction dynamics, which are essential for comprehending intracellular transport^32^, cannot be detected in raw WF images due to inadequate spatial resolution. UniSR overcomes this limitation by enabling the direct visualization of organelle motion and the tracking of interactions at high spatiotemporal resolution. The results in Figure 3 demonstrate that our method can resolve not only the morphologies of various subcellular structures but also their relative positions and motion trajectories, thus facilitating the study of interactions between different organelles during dynamic process. The enhancements in both contrast and resolution are stable during time-lapse network outputs, suggesting that UniSR can offer reliable monitoring of organelle interactions from indistinct captures.

### UniSR-enabled super-resolution imaging: resolution enhancement from confocal to STED-level images

To illustrate the generalization ability of our method across various imaging modalities and sample types, we implement the framework to diffraction-limited confocal images and present the outcomes in Figure 4. Fixed-cell NPCs, Golgi apparatus, and MTs, along with live-cell lysosomes and IMM, are imaged via an Abberior STEDYCON system (Methods). OMM in fixed cells are immunolabeled with anti-TOMM20 and acquired using our custom-built confocal/STED system^33^. Considering the specific challenges in staining lysosomes and IMM in fixed cells or obtaining high-quality, well-registered confocal-STED image pairs for these structures in live-cells, we formulate a degradation model. This model only requires SR images, which are computationally degraded to generate corresponding LR data (Methods), thus facilitating the convenient preparation of LR-SR image pairs for model fine-tuning. Figure 4A indicates that UniSR learns the mapping from diffraction-limited confocal images to STED SR images, thereby circumventing the use of complex optical setups and high-intensity depletion lasers to narrow the emission profile.

**Figure 4.**
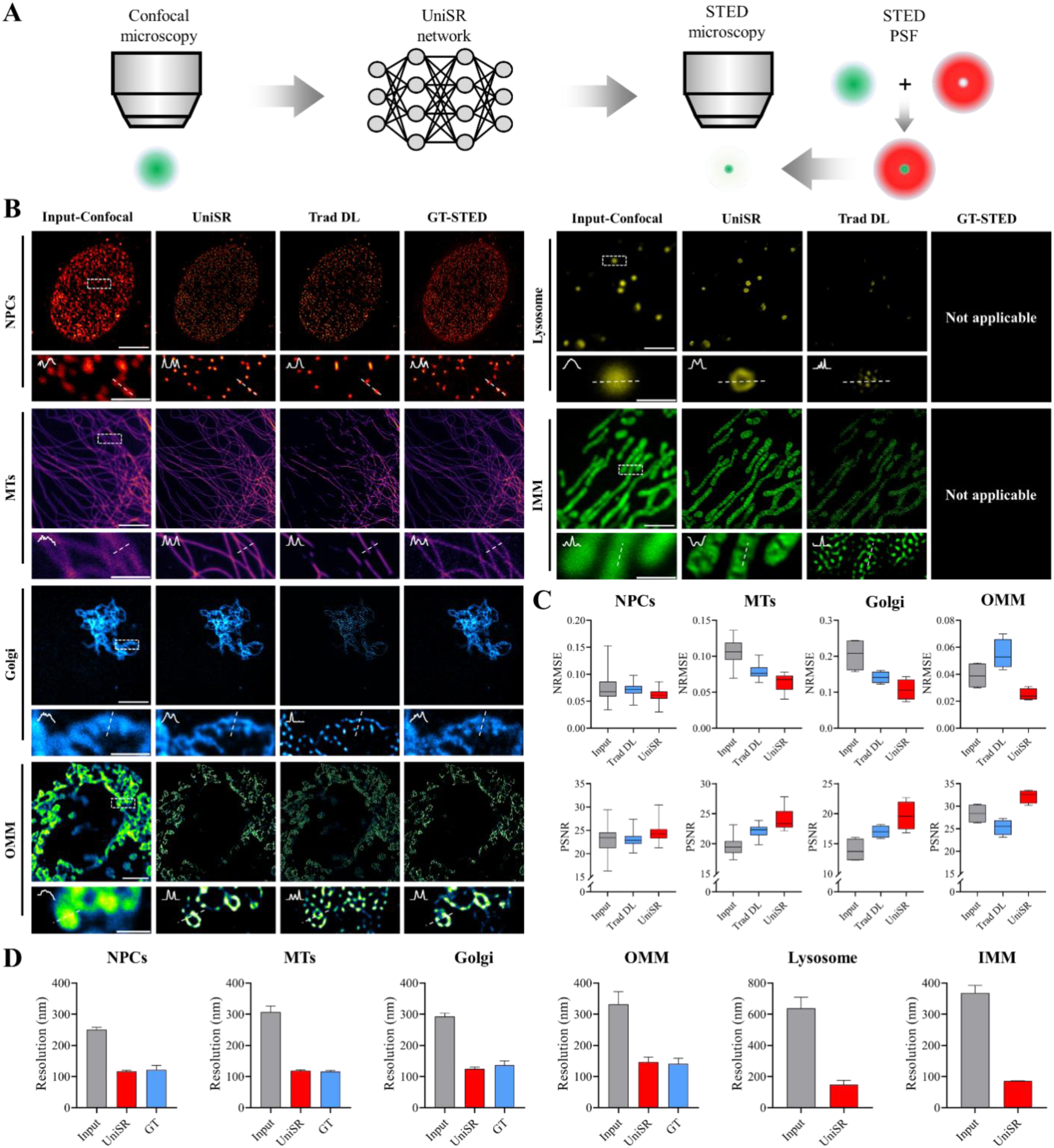
UniSR achieves STED-level resolution from confocal images. **A** Schematic representation of resolution improvement from confocal to STED through UniSR. **B** In the first column are diffraction-limited confocal images, which serve as inputs to the network. The second and third columns present the inferred SR images generated using UniSR and Trad DL, respectively. The fourth column displays the corresponding STED images of the same ROI. Magnified views of the areas within the white dashed boxes are presented below. **C**, **D** Comparison of NRMSE, PSNR, and resolution among the input confocal images, the inferred images obtained using UniSR and Trad DL, and the GTs. Scale bars: 5 μm (**B**), 1.25 μm (zoom-in regions in **B**).

UniSR persistently generates high-quality SR images of various organelles, displaying strong robustness when processing complex subcellular structures with high-fidelity. As shown in the magnified views of the white boxes in Figure 4B, the SR outputs of UniSR (the second column) present clearer and more comprehensive structures in comparison to the original confocal images (the first column) and the outputs of Trad DL networks (the third column). For example, the dense microtubules in UniSR predictions appear sharper and are more clearly resolved, and the OMM shows enhanced continuity. Moreover, the lysosomes and IMM in our results exhibit improved structural fidelity, while they are blurred in diffraction-limited confocal images or appear as fragmented, biologically implausible structures in traditional DL outputs. These findings are supported by the intensity profiles along white dashed lines: UniSR effectively suppresses artifacts and noise between adjacent structures observed in confocal images and overcomes the topological discontinuities commonly found in traditional DL results, thus preserving accurate cellular information. Notably, quantified by resolution measurements (Figure 4D), UniSR achieves resolution comparable to STED microscopy. Additionally, it is found that both NRMSE and PSNR of UniSR outputs are significantly better than those generated by Trad DL methods (Figure 4C).

SR images enable more precise quantitative analysis and subsequent biomedical research. For example, Figures. S9 and S10 present a comparison of the segmentation results of original LR images (WF or confocal), UniSR outputs, Trad DL outputs, and the GT (SIM or STED images) through the application of the minimum cross-entropy thresholding method^34^. Our proposed method enables the extraction of substantially more accurate morphological information compared to raw LR data or conventional deep neural networks. For instance, in the segmentation of UniSR output, the annular structures of the OMM network are clearly discernible, whereas they are scarcely distinguishable in the segmentation of both the input image and traditional DL output, which result in fragmented structures and false positives (Figure S10). This improved precision is of great significance for quantifying organelle size and density, parameters that exert a critical influence on cellular functions^35–37^. Moreover, UniSR further enhances the resolution when combined with deconvolution methods such as ZS-DeconvNet^10^ or Huygens (Figure S11).

## 3D UniSR-enabled volumetric super-resolution imaging

Three-dimensional (3D) imaging offers more comprehensive biological information compared to 2D observations. Nevertheless, it generally requires higher excitation light intensity and longer acquisition durations. This makes it more prone to phototoxicity, photobleaching, and out-of-focus background fluorescence ^10^. To overcome these limitations, we utilize the rapid adaptation capability and excellent SR performance of UniSR. We extend it to volumetric imaging by upgrading the network backbone to a 3D architecture (Figure S12A).

The 3D UniSR model is initially evaluated using public 3D datasets of NPCs and MTs^38^. It can effectively reconstruct fine structures from blurred LR images with high fidelity, achieving a resolution improvement of 2∼3 times, as quantitatively shown in Figure 5A. Although the Trad DL method can successfully perform SR on isolated MTs and circular NPC structures, it shows merging or loss of densely distributed targets. In contrast, UniSR can accurately perform SR on most structures without loss or merging, even when they are in close proximity to each other. This demonstrates significantly superior performance compared to conventional deep learning in volumetric SR inference for various biological specimens.

**Figure 5.**
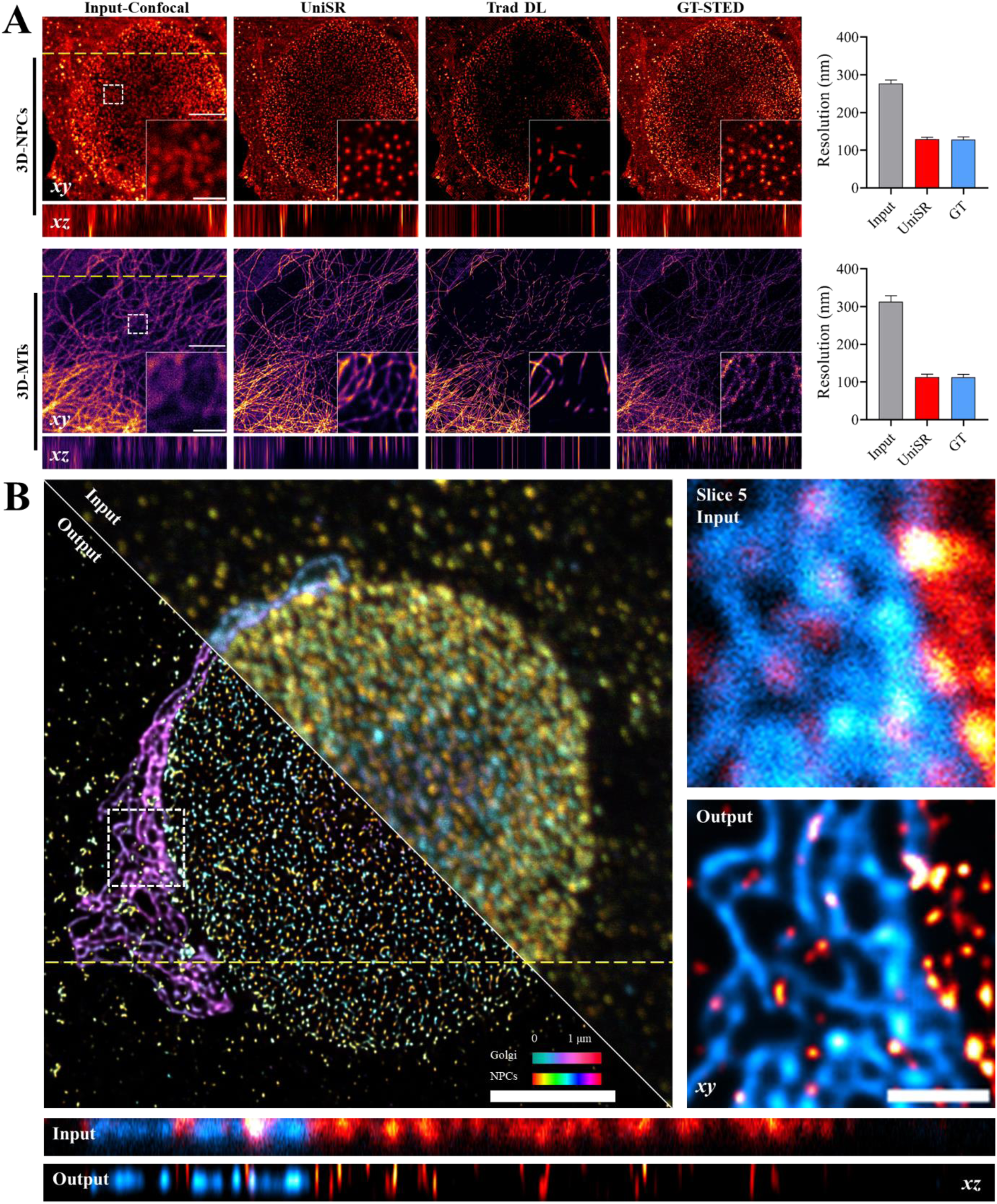
3D super-resolution imaging achieved by UniSR. **A** The first column presents 3D confocal volumes that serve as the input for the network. The second and third columns respectively display the inferred SR results obtained using 3D UniSR and Trad DL methods. The fourth column shows the corresponding STED volumes of the same ROI. Axial slices along the yellow dashed lines are presented below. Magnified views of white boxed regions are shown at the bottom right. The fifth column provides a quantitative comparison of the resolution among the input, UniSR outputs, and the GTs. **B** In the first column, the top-right part shows a 3D dual-color confocal dataset of Golgi and NPCs used as the input for the network, while the bottom-left part shows the 3D UniSR output. The second column displays magnified views of the regions within the white dashed boxes of the input (top) and the UniSR output (bottom). Axial slices of the input and the UniSR output are presented below. Scale bars: 5 μm (**A**), 2 μm (**B**), 1 μm (zoom-in regions in **A**, **B**).

To further assess the performance and extensive applicability of 3D UniSR, it is applied to a dual-color volumetric dataset of NPCs and Golgi proteins. This dataset is obtained using a ready-to-image slide (IG2COLOR-4006, abberior GmbH, Göttingen, Germany) and imaged with the commercial STEDYCON system. As depicted in Figure 5B, 3D UniSR significantly improves the SNR, contrast, and resolution of confocal volumes. It clearly resolves fine structural details that are indistinct in raw confocal images. The method also effectively captures the 3D distributions of Golgi and nuclear proteins, and their localization is clearly separated from the background noise. This facilitates the clear differentiation of the two organelles. In contrast, confocal images acquired under the same photon budgets are restricted by diffraction-limited resolution.

Moreover, it merits emphasis that the 3D UniSR model surpasses the 2D model in both the quality of the output images and spatial consistency (Figure S12B). Specifically, the former maintains the volumetric structural integrity across all three dimensions, whereas the latter gives rise to noticeable distortions along the axial dimension. This improvement is predominantly ascribed to the 3D UniSR’s capacity to comprehensively leverage global information across the entire volume. Conversely, the 2D model fails to efficiently collect cross-slice contents, leading to poor consistency between adjacent slices along the axial direction.

## Discussion

In this paper, we introduce UniSR, an algorithm based on transfer learning that is devised to directly generate SR images from commonly obtained diffraction-limited WF and confocal images. UniSR achieves SR capabilities comparable to conventional SR imaging techniques such as SIM, SMLM, and STED microscopy, while overcoming their key limitations. We illustrate its superior performance across a diverse range of organelles in multiple cell types, as elaborated in the Results section. Specifically, these encompass MTs in COS-7, Beas2B, HeLa, U2OS, and mouse embryonic fibroblasts; lysosomes in COS-7 and HeLa cells; OMM in COS-7 and HeLa cells; F-actin in COS-7 cells; IMM in HeLa cells; CCPs in COS-7 cells; ER in COS-7 cells; NPCs in HeLa, Vero, and mouse embryonic fibroblasts; and Golgi apparatus in Vero cells. Beyond achieving SR of various cellular structures and dynamics, UniSR demonstrates robustness against alterations in imaging conditions including eight different microscopy setups, SNR levels, FOVs, and both 2D and 3D imaging, which underscores its generalizability and wide-ranging applicability.

The proposed methodology constructs a robust framework that integrates SR capabilities with the advantages of WF or confocal microscopy, thereby surmounting the intrinsic trade-offs among spatial resolution, imaging velocity, and light dosage in conventional SR systems. These benefits are of particular significance for live-cell imaging, especially during the long-term surveillance of organelle dynamics and interactions when the SNR is exceedingly low. UniSR computationally enhances the resolution of structural details and generates high-fidelity images under diverse noise levels (Figure 4 and Figure S3). It is also capable of achieving SR from a single WF image, significantly mitigating specimen photobleaching (Figures. 2 and 3). This effectively addresses a crucial limitation of conventional SR imaging, which generally requires relatively high excitation laser power to acquire an adequate fluorescence signal for visualizing delicate subcellular structures, unavoidably leading to phototoxicity, photobleaching, and other adverse effects. By obviating this requirement, UniSR enables long-term observations of biological processes without inflicting discernible photodamage on cells.

This study demonstrates the robust generalization ability of UniSR, which effectively transfers representations from a pre-trained model to specific application tasks. Through efficient fine-tuning with only one image pair (Figure S8), UniSR can rapidly adapt to perform SR on various subcellular structures. This approach circumvents the necessity for other deep-learning-based methods to collect extensive high-quality training data, rendering SR imaging feasible for biological structures where the acquisition of a large volume of training datasets is laborious or challenging. We pre-train UniSR using simulated data and achieve performance comparable to that of the same model pre-trained on experimental data (Figure S7). This approach presents significant practical advantages, as the generation of simulated data is straightforward (Figure S13), thus saving time on experimental data acquisition. More crucially, when the imaging modality or specimen type change, users can directly fine-tune the pre-trained model instead of retraining it from scratch, resulting in a reduction of training time. To enhance the accessibility of UniSR, we develop a Fiji^39^ plugin toolbox (Video S1). This toolbox is user-friendly, enabling even those without deep learning experience to fine-tune the models.

UniSR presents a novel multi-window synchronized sampling strategy (Methods) to comprehensively capture feature information from a single fine-tuning image pair, thus enhancing learning efficiency. Additionally, a multi-component loss function is designed, which optimizes both the learning process and model performance. This leads to the generation of SR structural details and enables more distinct visualization of subcellular morphology, organelle dynamics, and interactions. Quantitative assessments employing NRMSE, PSNR, and SSIM metrics demonstrate substantial improvements in image quality and resolution of the UniSR outputs, when compared to both LR inputs and the outcomes of Trad DL method. Intensity profiles further display significantly improved visibility and structural consistency with GT images acquired through conventional SR techniques. Moreover, UniSR is extended to a 3D model capable of super-resolving fine subcellular structures (e.g., NPCs, MTs, Golgi apparatus) from diffraction-limited confocal volumes. This facilitates rapid and high-fidelity volumetric SR observation of multiple organelles (Figure 5).

These findings indicate that the outstanding SR capability of UniSR paves the way for more in-depth exploration of subcellular dynamics. It positions itself as a powerful computational SR microscopy tool, especially for the study of biological processes that pose challenges to traditional SR imaging techniques. As a universal, data-driven methodology, UniSR requires no alterations to existing WF or confocal systems. Additionally, it can be employed to improve the resolution in other imaging modalities, such as electron microscopy (EM) (Figure S14).

In summary, UniSR establishes a broadly applicable computational framework for subcellular SR fluorescence imaging. By combining physics-informed pre-training with minimal fine-tuning, it achieves robust and generalizable performance across diverse imaging modalities, noise levels, and biological specimens. These capabilities make UniSR a practical tool for live-cell studies and point toward new opportunities to integrate artificial intelligence with advanced microscopy for biological discovery.

## Methods

### UniSR framework

#### UniSR core

The UniSR framework constructs its computational basis via a novel amalgamation of cross-modal feature alignment and transfer learning. The former emphasizes minimizing distributional disparities between source and target images, whereas the latter utilizes pre-aligned features to enhance image resolution through fine-tuning. Notably, this approach exhibits remarkable efficiency by attaining significant resolution improvement with one pair of LR and SR training data. Although the transmission of low-frequency information is independent of transfer learning, our findings indicate that the strategic application of transfer learning promotes the retention of crucial high-frequency components that are essential for resolution enhancement.

Conventional transfer learning methods for SR generally require substantial pre-training on large paired datasets obtained from the target imaging system and its corresponding SR system to establish prior knowledge of resolution mapping. This data-intensive strategy is fundamentally at odds with our goal of data efficiency. To tackle the problem, we generate simulated structures for pre-training the model. A crucial requirement is to maintain a consistent resolution scaling factor between simulated LR and SR image pairs, while allowing flexibility in other structural parameters. The subsequent fine-tuning only requires one pair of experimental LR-SR images to adjust the model weights and achieve mapping refinement. As a result, UniSR can effectively enhance the resolution while reducing the demand for real data compared to conventional approaches.

#### Multi-window sampling strategy

Considering the fine-tuning with only one LR-SR image pair, maximizing information utilization is crucial for improving the generalization capacity of the UniSR. To achieve this objective, we adopt a multi-window sampling strategy. This strategy randomly extracts eight 256 × 256 patches from each image, thereby increasing the input data and achieving feature representation with diverse spatial coverage across the entire image. The patch number is constrained by GPU memory, though in theory, more patches would further increase feature diversity. During the training iterations, the model progressively extracts and leverages information from the data pair, thereby enhancing its learning ability.

To augment dataset diversity via geometric augmentation, we employ a stochastic transformation in which each input image has a 50% probability of remaining unchanged. When transformations are applied (with a 50% probability), we randomly select from seven distinct spatial operations in a uniform manner: (1) vertical flipping, (2) 90° clockwise rotation, (3) 90° clockwise rotation followed by vertical flipping, (4) 90° counter-clockwise rotation, (5) 90° counter-clockwise rotation followed by vertical flipping, (6) 180° rotation, and (7) 180° rotation followed by vertical flipping.

The multi-window sampling strategy facilitates the concurrent acquisition of information from various locations. It retains high-frequency components within local regions and improves structural accuracy by leveraging global contextual relationships among image patches. Through the implementation of this strategy, UniSR achieves reliable feature representation even with limited training sample, thereby enhancing the quality of SR output.

#### Network architecture

We develop a dual-attention network (DA-net) similar to our previous work in Ref.^40^. This network is founded upon the encoder-decoder framework and draws inspiration from U-net ^41^, the residual channel attention network^42^, and Transformer^43^. The encoder extracts spatial features via four downsampling blocks, each of which encompasses two residual channel attention blocks (RCAB) and a convolutional layer. The channel attention (CA) within RCAB enhances prominent structures, retains crucial details, and bolsters the network’s capacity to capture high-frequency information. Let *x* be the input of RCAB, and the feature maps in each RCAB are

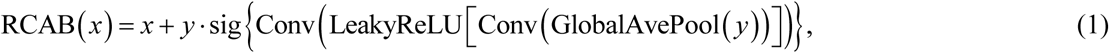

where

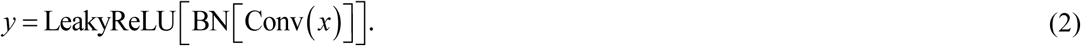

In equation (1), the channel-wise global spatial features are compressed through global average pooling GlobalAvePool(), as described in Ref.^42^. We implement two convolution operations Conv() with a kernel size of 1 × 1 and a stride of 1 for feature extraction. A leaky rectified linear unit activation function LeakyReLU[] with a slope of *α* = 0.2 is employed to introduce nonlinearity^44^. Mathematically, LeakyReLU (*x*_in_; *α*) = max(0, *x*_in_) − *α* · max(0, −*x*_in_). Subsequently, a sigmoid activation sig{} normalizes the attention weights^45^. In equation (2), a convolution operation with kernel size of 3 × 3, a stride of 1, and a padding of 1 is utilized to enhance model accuracy, followed by batch normalization BN[] to mitigate internal covariate shift and expedite convergence^46^.

The decoder converts the feature maps into SR images via four upsampling blocks. Each of these blocks comprises two RCABs and one deconvolutional layer. The skip connections concatenate the features from the corresponding encoder and decoder layers. Subsequent to the final upsampling block, a deconvolutional layer with a kernel size of 2 × 2 maps 64 channels into 1 channel to generate a high-quality grayscale image with the same pixel dimensions as the input images.

A self-attention (SA) module is incorporated into the transition layer, enabling the model to capture long-range dependencies and structural details^47^. It calculates the query, key, and value matrices for a local window feature and implements an attention mechanism. These matrices are normalized within group channels^48^ and refined through a convolution layer, guaranteeing training stability. The multi-head attention module employs eight parallel heads. Each head applies relative positional encoding with learnable parameters to the attention scores prior to softmax normalization, after which the head outputs are concatenated.

To perform SR on 3D data (xyz coordinate system), we devise a volumetric extension of DA-net to comprehensively utilize axial spatial information. All 2D operators are enhanced to native 3D implementations. For instance, 3D kernels and voxel-wise attention are employed instead of 2D spatial convolutions and attention respectively, thus establishing a 3D-adapted SR model for volumetric data processing.

#### Loss function

To precisely perform SR of subcellular structures, the loss function is designed as a combination of mean square error (MSE), SSIM, and feature reconstruction loss (FRL). Parameters *α*, *β* and *γ* are employed to balance the corresponding terms. In our experiments, *α* = 0.8, *β* = 0.2, and *γ* = 0.05 are empirically selected to optimize performance, resulting in the following formulation:

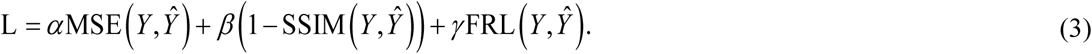

The MSE loss guarantees prediction accuracy by penalizing the discrepancy between the network output 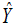 and ground truth *Y*, while the SSIM loss improves the perceptual quality of the output. The FRL is integrated to enforce feature-level alignment between the output image and corresponding target, which is given by

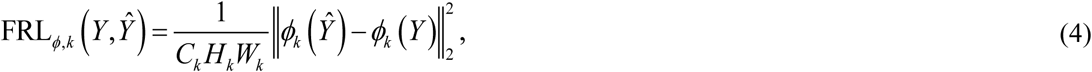

where *ϕ_k_* () is the activation of the *k*th layer of the network *ϕ*. If *k* is a convolutional layer, then *ϕ_k_* () will be a feature map of shape *C_k_* × *H_k_* ×*W_k_* ^49^. Here, the 30th layer in the VGG-16 network^50^ is adopted as our *ϕ_k_* () to extract high-level features and promote the output image to be perceptually similar to the target image, rather than precisely matching its pixels.

#### Simulated pre-training data

UniSR is pre-trained exclusively on simulated data, and the generation of the simulated LR-SR image pair is adapted from Ref.^15^. Taking the line structure as an example (Figure S13), initially, 3 to 50 lines with random orientations are sketched on a blank image of 20 × 20 μm^2^ with a pixel size of 40 nm. Subsequently, a Gaussian mask of *σ* = 2 pixels is utilized to randomly disrupt the continuity of the line structures, thereby introducing random notches along the lines. Next, elastic deformation (two typical deformation fields are presented in Figure S13) is applied to transform the straight lines into curved lines. Finally, the resulting image is convolved with Gaussian kernels having full widths at half maximum (FWHM) of 130 nm and 380 nm to generate the simulated SR and LR images, respectively. Analogously, simulated images of circular and annular structures are generated in accordance with these procedures, with each image containing a random distribution of 150 to 300 circles and annuli.

#### Training

The training process consists of two stages: pre-training and fine-tuning, which utilize prior knowledge to improve the model’s adaptability to new tasks. In the pre-training stage, a total of 5000 simulated images with a size of 256 × 256 pixels are generated. Subsequently, these images are partitioned at a ratio of 4:1 for training and validation purposes. Approximately, 14 hours are required to complete 500 epochs for pre-training, with an initial learning rate of 1 × 10^-4^ and a batch size of 8. During the fine-tuning stage, the pre-trained network is optimized on a modality-specific dataset. All layers except the last convolutional layer were frozen, restricting parameter updates to the weights of that layer. Approximate 10 minutes are needed to complete 500 epochs for fine-tuning, with an initial learning rate of 8 × 10^-5^, which suffices to achieve a SR image. The required training time for fine-tuning is primarily determined by the imaging modality and the structural complexity of cellular organelles. 3D volumes require significantly longer optimization periods compared to conventional 2D slices.

We conduct min-max normalization on both input and target images, normalizing pixel intensities to the interval of [0,1]. This preprocessing step ensures numerical stability in gradient computations while preserving relative intensity relationships among different images.

The training procedure employs the adaptive moment estimation (Adam) optimizer^51^ in conjunction with a cosine annealing learning rate scheduler. This computational framework is constructed using Pytorch 1.13.0^52^ and Python 3.8.3 under Microsoft Windows 10 operating system. All training and inference operations are carried out on a workstation furnished with an NVIDIA GeForce RTX 3080 GPU and an AMD Ryzen 5 5600X CPU @3.7 GHz.

### Performance evaluations

#### Pixel-wise metrics

In this work, NRMSE, SSIM and PSNR are employed as metrics to evaluate the pixel-level consistency between network output and GT images. Their respective calculations are as follows:

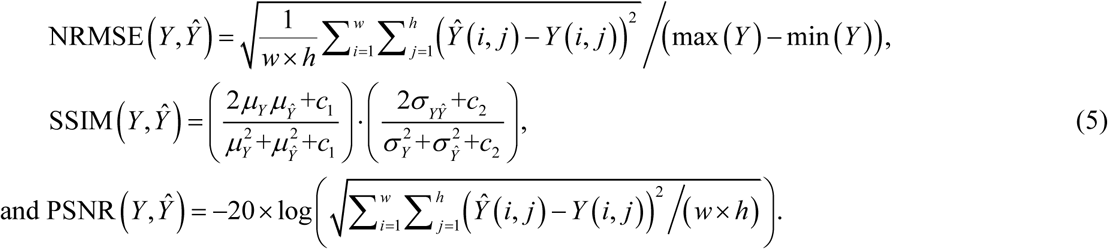

where the width and height of the images are denoted by *w* and *h*, respectively. *μ* and *σ* represent the mean and standard deviation of the images, respectively, and 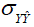 is the covariance of the network output 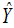 and the ground-truth image *Y*. Both *c*_1_ and *c*_2_ are small constants introduced to ensure the stability of the division.

#### Image resolution estimation

Image resolution is achieved through decorrelation analysis, which characterizes the highest frequency based on the local maxima of the decorrelation functions rather than the theoretical resolution^28^. We utilize its ImageJ plugin while configuring the pixel size and other default parameters.

### WF-SMLM image acquisition

#### Cell preparation

COS-7 cells are treated by paraformaldehyde (PFA)-glutaraldehyde (GA) mixed fixatives (4% PFA + 0.1% GA) for 30 minutes at room temperature. Then, cells are permeabilized and blocked in a blocking buffer consisting of 3% (w/v) BSA and 0.2% (v/v) Triton X-100 in PBS for 20 minutes. α-tubulin primary antibody incubation is performed in blocking buffer for 1 hour. After three washes with washing buffer (0.2% BSA, 0.1% Triton X-100 in PBS), samples are incubated with secondary antibody for 1 hour at room temperature. Finally, cells are washed three additional times before SMLM imaging. PFA-GA mixed fixatives (NWFix#2), mouse monoclonal anti-α-tubulin antibodies (NW-Tubulin#1-M) and Alexa Fluor 647-conjugated anti-mouse IgG (H+L) secondary antibodies (NWSM647#1) are obtained from Nano-Microimaging Biotechnology (Ningbo, China).

#### Optical setup and data acquisition

The cells on the 12-mm coverslips are mounted on 22 mm × 60 mm glass slides for WF and SMLM SR imaging. WF images are directly collected by an inverted fluorescence microscope (Ti2-E, Nikon, Japan). SMLM images are obtained by a commercial SMLM setup (STORM Ultra300, Nano-Microimaging Biotechnology, Ningbo, China) based on an inverted fluorescence microscope (Ti2-E, Nikon, Japan) equipped with a sCMOS camera (Dhyana 95V2, TUCSEN, Fujian, China) and a 100× oil objective (Nikon CFI Plan Apochromat λ, NA 1.49). A 647 nm laser illuminates the sample at ∼2 kW/cm^2^ to photoswitch most of the labeled AF647 dye into a dark state. A weak 405 nm laser (typically 0∼1 W/cm^2^) is used to stochastically reactivate fluorophores into the activation state to allow single-molecule fluorescence detection. The single-molecule images are obtained at ∼100 frames per second. Typically, SMLM images are reconstructed from ∼50,000 frames according to localization algorithm.

### 2D confocal-STED image acquisition

#### Cell preparation

For fixed cell, they are fixed with 4% paraformaldehyde (Sigma-Aldrich, P6148) at room temperature for 30 minutes. After fixation, cells are permeabilized with 0.5% Triton X-100 for 30 minutes and blocked in 2% BSA for 1 hour. Primary antibody incubation (e.g., anti-TOMM20, Abcam EPR15581-54) is performed overnight at 4°C. Secondary antibodies conjugated to fluorophores are applied at room temperature for 1 hour prior to mounting and imaging. For live cells, they are fixed at 37°C using a prewarmed fixation solution that contains 4% paraformaldehyde and 0.1% glutaraldehyde (SunBloss™, HXKx01, prepared in PBS) for 10 min. After fixation, the samples are washed three times with PBS. To reduce background fluorescence, the cells are gently shaken in a PBS solution containing 0.1% NaBH₄ (SunBloss™, HXIK023) for 7 min (< 1 Hz), followed by three washes with 2 ml PBS. The cells are then incubated at 37°C for 30 min in a blocking solution that is composed of PBS with 5% BSA and 0.5% Triton X-100 (SunBloss™, HXKx02). The primary antibodies are diluted in the same blocking solution. The samples are incubated at 25°C for 40 min with the following primary antibodies: β-tubulin (DSHB-E7) and Tom20 (ABclonal, A19403). After incubation, the samples are washed three times with 2 ml PBS for 5 min each. Subsequently, the samples are incubated at 25°C for 60 min with secondary antibodies (Goat anti-mouse IgG, STRED-1001-500UG; Goat anti-rabbit IgG, STRED-1002-500UG; both from Abberior), while protected from light. After incubation, the samples are washed three times with PBS and post-fixed for 10 min. For live-cell samples, 2 × 10⁵ cells are seeded in glass-bottom microwell dishes and cultured in 1 ml DMEM containing 10% FBS for 24 h. After overnight incubation, the cells are washed three times with PBS.

#### Optical setup and data acquisition

The STEDYCON system (Abberior) is built on an inverted microscope (Olympus) equipped with a 100×/1.45 oil immersion objective (HCX PL APO, 100×/1.40-0.9 OIL, Leica, Germany). Excitation is achieved through 640 nm laser illumination, while depletion is accomplished with a 775 nm laser. To prevent spectral overlap and maintain image resolution, organelle-specific staining is performed using spectrally distinct fluorophores. NPCs within fixed cells (stained with STAR Red) are imaged with 5-line accumulation and 30 nm pixel resolution. Microtubules within fixed cells (stained with STAR Red) are imaged with 5-line accumulation and 30 nm pixel resolution. Golgi is from commercially available fixed samples of Abberior, with 30 nm pixel resolution. Lysosomes within live MDA cells (stained with LysoBrite^TM^ NIR) are incubated for 15 minutes and imaged with 3-line accumulation and 30 nm pixel resolution. IMM within live Hela and MDA cells (stained with PK Mito Orange) are incubated for 15 minutes and imaged with 3-line accumulation and 20 nm pixel resolution.

#### Generation of synthetic confocal images

Inspired by Ref.^38^, a degradation model is developed to generate LR data when we can only collect SR STED images. This allows us to prepare conveniently well-registered image pairs to fine-tune the network. As illustrated in Figure S15, if the confocal image *I*_C_ and STED image *I*_STED_ are regarded as the convolution of the object *obj* with their respective system point spread function (PSF), then we formulate the imaging process as

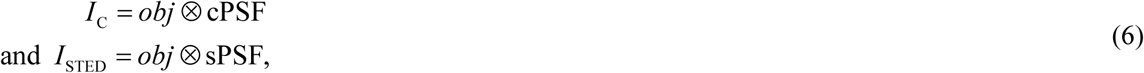

where ⊗ denotes the convolution operation. cPSF and sPSF are estimated PSF of confocal and STED imaging system, respectively. In the first step, by applying Fourier transform FFT () to equation (6) and dividing the first equation by the second one, we eliminate the object spectrum and obtain

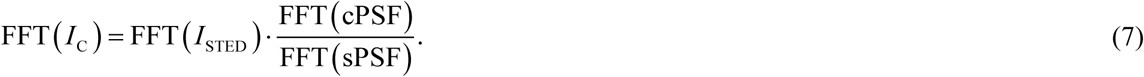

In the next step, inverse Fourier transform iFFT() is applied to the above equation to yield a synthetic confocal image, and it is the degraded LR data from the corresponding high-resolution STED image. In the third step, the synthetic image is further corrupted with Gaussian and Poisson noise to better resemble experimentally acquired LR image.

### Open-sourced datasets

#### WF-SIM images

We use the public dataset BioSR dataset from Ref.^15^ and BioTISR dataset from Ref. ^14^ to evaluate our UniSR’s resolution enhancement performance on WF images. We apply bilinear interpolation to upsample WF images by a factor of two due to the inherent pixel resolution mismatch between WF and SIM images. This computational alignment strategy achieves inter-modality dimensional consistency while minimizing interpolation-induced artifacts.

#### WF-SMLM images

We use the public dataset from Ref.^16^ to test our UniSR’s resolution enhancement performance on WF images (Figures. 3C and 3D).

#### 3D confocal-STED images

We use the public dataset from Ref.^38^ to test our UniSR’s resolution enhancement performance on 3D confocal images (Figure 5A).

#### EM images

We use the public dataset from Ref.^53^ to test our UniSR’s resolution enhancement performance on EM images (Figure S14).

## Data Availability

The data that support the findings of this study are available from the corresponding author upon reasonable request.

## Code Availability

The codes that support the findings of this study are openly available at https://github.com/Xu-onion/UniSR.

## Acknowledgements (optional)

This work is supported by the National Key Research and Development Program of China (2021YFF0502900); the National Natural Science Foundation of China (T2421003, 62127819, 62435008, 62435011); the Natural Science Foundation of Guangdong Province (2024A1515012082).

## Supplementary Information

Figures. S1-S15

Video S1. UniSR tutorial for rapid application in the Fiji plugin toolbox (related to Figure 1).

